# AGEing of Collagen: The Effects of Glycation on Collagen’s Stability, Mechanics and Assembly

**DOI:** 10.1101/2024.07.09.602794

**Authors:** Daniel Sloseris, Nancy R. Forde

## Abstract

Advanced Glycation End Products (AGEs) are the end result of the irreversible, non-enzymatic glycation of proteins by reducing sugars. These chemical modifications accumulate with age and have been associated with various age-related and diabetic complications. AGEs predominantly accumulate on proteins with slow turnover rates, of which collagen is a prime example. Glycation has been associated with tissue stiffening and reduced collagen fibril remodelling. In this study, we investigate the effects of glycation on the stability of type I collagen, its molecular-level mechanics and its ability to perform its physiological role of self-assembly. Collagen AGEing is induced *in vitro* by incubation with ribose. We confirm and assess glycation using fluorescence measurements and changes in collagen’s electrophoretic mobility. Susceptibility to trypsin digestion and circular dichroism (CD) spectroscopy are used to probe changes in collagen’s triple helical stability, revealing decreased stability due to glycation. Atomic Force Microscopy (AFM) imaging is used to quantify how AGEing affects collagen flexibility, where we find molecular-scale stiffening. Finally we use microscopy to show that glycated collagen molecules are unable to self-assemble into fibrils. These findings shed light on the molecular mechanisms underlying AGE-induced tissue changes, offering insight into how glycation modifies protein structure and stability.

## INTRODUCTION

Aging is a multifaceted biological process characterized by the progressive decline of physiological functions and an increased vulnerability to a range of diseases [1, 2]. While many mechanisms underlie aging, a key molecular pathway involves non-enzymatic glycation, which results in the formation of a variety of age-related Advanced Glycation End-products (AGEs) [3]. Although select AGEs have been shown to be removable, they are in general considered to be irreversible once fully formed [4]. AGEs have been linked to progression of cardiovascular disease [5, 6], cancer metastasis [7–9], tissue stiffening [10, 11] and increased rate of injury and degeneration [3, 12]. Further, AGEs have been shown to be useful as diagnostic biomarkers for age-linked diseases [3, 12]. Although glycation occurs constitutively, AGEs accumulate slowly and therefore mostly on proteins with slow turnover rates such as collagen [10]. The chronic hyperglycemia associated with diabetes significantly accelerates and exacerbates glycation’s harmful effects, leading to more rapid accumulation of AGEs [3, 10]. This accelerated glycation process in diabetics contributes to the deterioration of collagen’s structural integrity and mechanical properties. As collagen mechanics influence cellular function, the effects of chemical composition on collagen mechanics and stability at the molecular level are pivotal in understanding age-related changes in the human body [13].

Collagen is a triple-helical protein found in the extracellular matrix that provides support, elasticity and stability to tissues [13]. At the molecular level, collagen is a ∼300 kDa protein, structured as a ∼300 nm long right-handed triple helix ∼1.5 nm in diameter, comprising 3 left-handed helical *α*-chains [13]. Type I collagen, the most abundant type in the body, is one of the fibril-forming collagens, meaning that in the body it self-assembles into higher-order fibrillar structures (Fig. 1A).

**FIG. 1.**
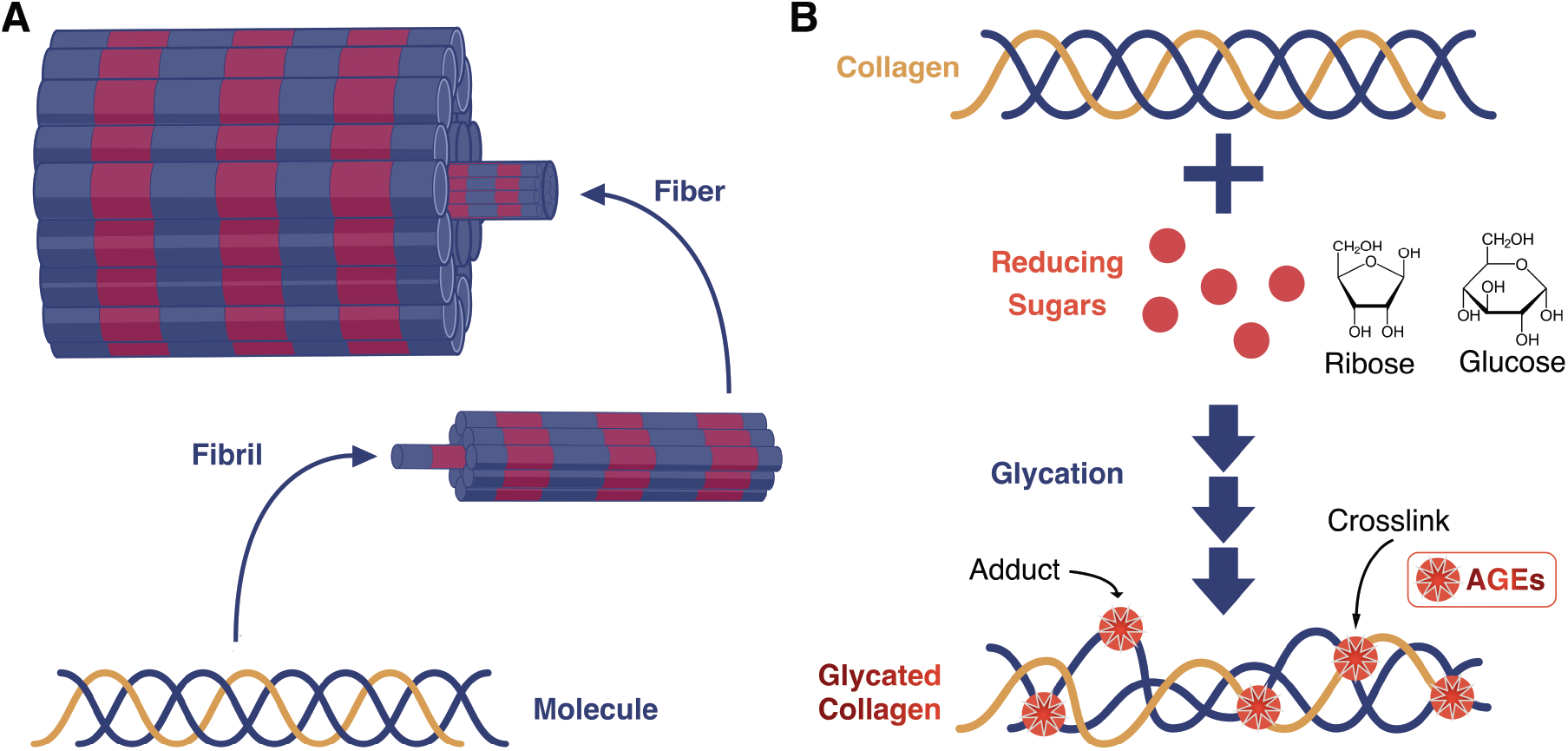
Collagen structure and glycation. (A) Visual representation of different structural levels of collagen type I organization. Banding is meant to represent gap and overlap regions found in assembled fibrils and is not to scale. Type I collagen is comprised of two identical *α*1 (blue) chains and one *α*2 (yellow) chain. (B) Glycation of collagen. Sugars including ribose and glucose can participate in glycation. Reactive amino groups found on lysine and arginine residues commonly participate in AGE formation. AGEs can modify a single site to form an adduct or two adjacent sites to form a crosslink.

AGE formation proceeds via the Maillard reaction, the same reaction that browns bread and meat while cooking: reducing sugars (*in vivo*, predominantly glucose) react with amino groups of amino acids (typically lysine and arginine) to form crosslinks and adducts (Fig. 1B) [10]. Late-life collagens bear the marks of decades of accumulated AGEs, manifesting as intra-molecular and inter-molecular crosslinks between *α*-chains and adducts attached to a single *α*-chain (Fig. 1 B) [10]. These modifications result in stiffening of fibrils and tendons, and altered thermal stability and proteolytic susceptibility [10, 12, 14–16]. However, most of the literature has focused on identifying glycation-induced changes of collagen fibrils. Here we instead focus specifically on changes of individual collagen proteins in order to understand the effects of glycation of collagen at its most fundamental structural level.

In this study, we explore how glycation impacts the stability and flexibility of molecular type I collagen, as well as its ability to self-assemble post-glycation. We employ a multifaceted approach, combining traditional biochemical methods with single-molecule imaging and analysis techniques to unveil the mechanical alterations associated with advanced glycation end products (AGEs) at the level of individual proteins.

## EXPERIMENTAL PROCEDURES

### Glycation of Collagen

Collagen (Cultrex rat tail type I, 3440-100-01) was glycated in acidic conditions (20 mM acetic acid at pH = 3.2) at a concentration of 2 mg/mL with varying concentrations of D-ribose (Sigma, 50-69-1). All samples were incubated at room temperature for varying lengths of time from 7 - 350 days and then, unless otherwise noted, were dialyzed into 20 mM acetic acid and stored at 4°C. Control samples were treated identically, but omitting ribose.

### SDS-PAGE

Aliquots of collagen samples incubated at various D-Ribose concentrations were subjected to SDS-PAGE electrophoresis in a 6%/5% separating/stacking polyacrylamide gel using Bio-Rad (USA) electrophoresis equipment. Samples were denatured by boiling at 95°C prior to electrophoresis under non-reducing conditions.

### Fluorescence Measurements

Fluorescence readings were taken on a Tecan Infinite 200 Pro plate reader at room temperature. Wavelengths of *λ*_ex_ = 370 nm (excitation) and *λ*_em_ = 440 nm (emission) were used to monitor general AGE fluorescence [17]. Wavelengths of *λ*_ex_ = 328 nm and *λ*_ex_ = 378 nm were used to monitor pentosidine-specific fluorescence [18]. Fluorescence measurements were taken at a collagen concentrations of 0.1 mg/ml.

### Trypsin Digestion

Digestion of collagen samples was conducted at 37°C for 30 minutes in 50 mM Tris, 200 mM NaCl pH 7.5 with 0.25 mg/mL trypsin (Sigma) and 0.3 mg/mL collagen. After digestion, samples were subjected to SDS-PAGE. SDS-PAGE gels were analyzed using ImageJ. Percent digestion was computed by comparing integrated band intensity through a line scan for lanes containing collagen with and without trypsin. Integrated intensities were summed of both alpha bands or of both beta bands and compared to their undigested counterparts.

### Circular Dichroism (CD) Spectroscopy

CD measurements were recorded on a Chirascan V100 using a 1 mm quartz cuvette (Starna). Collagen samples were diluted into 20 mM acetic acid to a final concentration of 0.2 mg/mL. Baselines were collected using 20 mM acetic acid. Spectra were measured from 190 nm to 260 nm, with presented spectra representing an average of at least 5 measurements. These averaged spectra were then scaled by dividing by the maximum value of the ∼ 222 nm peak. Temperature ramps were recorded at 222 nm from 34°C - 50°C at a rate of 0.5°C per minute. Biphasic sigmoid curves (a combination of two logistic functions) were fit to the temperature-dependent amplitudes of these thermal denaturation curves using the following equation:

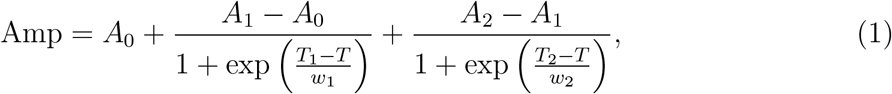

where the fitting parameters *A*_0_, *A*_1_, *A*_2_ are amplitudes, *T*_1_ and *T*_2_ are the midpoints of the respective transitions, and *w*_1_ and *w*_2_ are the widths of the transitions. Single-sigmoid models were also tested, in which the final term from this equation was omitted. To be agnostic to the choice of fitting model, melting temperatures *T*_m_ were identified as the temperature at which the amplitude was ½ of its initial value.

### Atomic Force Microscopy (AFM)

AFM imaging was performed as per [19], using Oxford Instruments Asylum Research MFP-3D and Jupiter AFM instruments, operating in tapping mode in air. Collagen samples were diluted into 1 mM HCl, 100 mM KCl (pH ∼3) at a final concentration of ∼1 *µ*g/mL; 50*µ*L of the diluted sample was deposited and allowed to settle for 30 s on freshly cleaved mica (Highest Grade V1 AFM Mica Discs, 10 mm; Ted Pella, Redding, CA). Unbound proteins were removed by rinsing with ultrapure water, and the mica was then dried using filtered air. All proteins were imaged under dry conditions.

Image analysis (tracing of collagen backbones and fitting to the Worm-Like-Chain (WLC) model) was performed using AutoSmarTrace and SmarTrace software [19, 20]. In short, collagen molecule backbones were traced using the automated tracing software AutoSmarTrace [20], and the traced backbones were then separated into segment lengths of *s* = 10 −200 nm. For each segment, the mean-squared end-to-end distance ⟨*R*^2^(*s*)⟩ and the tangent vector correlation ⟨cos *θ*(*s*)⟩ were determined and used to estimate persistence length *p* by comparing to the WLC predictions for equilibrated polymers in 2D [19]:

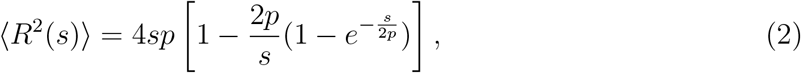

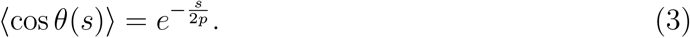

Error estimation of persistence length was performed using bootstrapping. All *N* traced chains were resampled into subsets containing *N* /2 chains, with repetition over 30 iterations. Persistence lengths determined via ⟨*R*^2^(*s*)⟩ and ⟨cos *θ*(*s*)⟩ were averaged for each bootstrapping iteration. The mean and standard deviation of these 30 bootstrapped persistence lengths are reported.

### Fibril Assembly

Differential Interference Contrast (DIC) microscopy was employed to visualize the assembly of collagen type I into fibrils, following neutralization in phosphate-buffered saline (PBS, pH = 6.9) and overnight incubation at room temperature [21, 22]. Samples were then pipetted onto microscopy slides and imaged using DIC microscopy at 40X magnification on an Olympus IX83 microscope.

To assess the impact of free ribose on fibril formation, turbidity was measured at 347 nm using a Synergy HTX plate reader.

## RESULTS

### Collagen is glycated by ribose in a concentration-dependent manner

In this study, we aimed to induce AGEs in collagen by incubating with ribose at different concentrations, and for different durations of time; and then to determine the effects of AGEs on collagen’s properties. First, we used fluorescence and SDS-PAGE to confirm that ribose incubation in acidic conditions – used to maintain collagen in soluble, molecular form – introduced AGEs into collagen.

Fluorescence readings at general AGE wavelengths (*λ*_ex_ = 370 nm, *λ*_em_ = 440 nm) [17] and at pentosidine-specific wavelengths (*λ*_ex_ = 328 nm, *λ*_ex_ = 378 nm) [18] showed more intensity at higher ribose concentrations, indicating greater AGE presence (Fig. 2A). Control samples incubated with 0 M ribose displayed fluorescence signals consistent with blanks (20 mM AcOH) across various measurements.

**FIG. 2.**
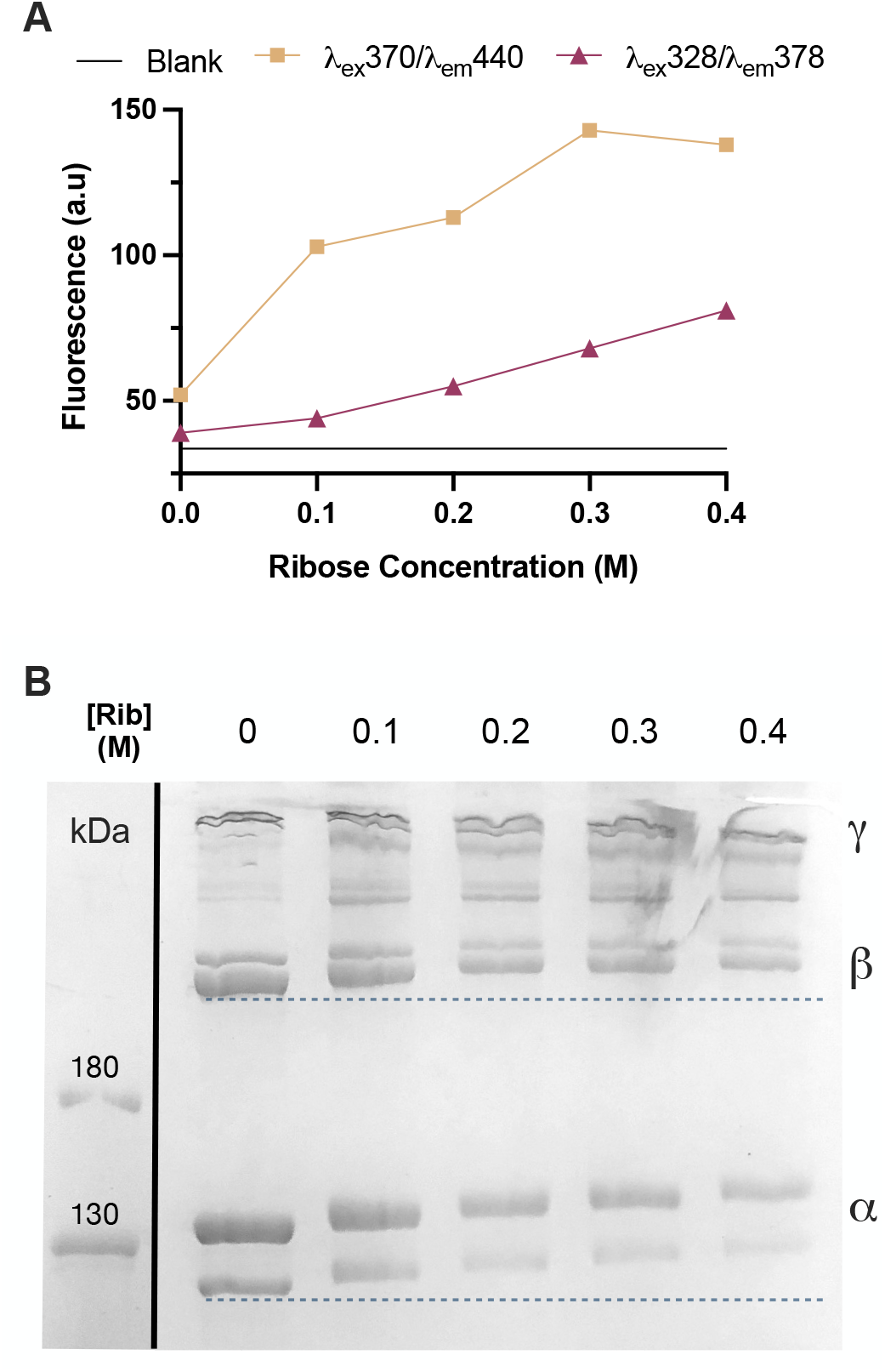
Confirmation and tracking of collagen glycation progression. (A) Fluorescence measured at *λ*_ex_ = 370 nm, *λ*_em_ = 440 nm was used to track general AGE presence while fluorescence at *λ*_ex_ = 328 nm, *λ*_em_ = 378 nm corresponds to pentosidine. Both signals increase with the concentration of ribose used for treatment. (B) Denaturing SDS-PAGE gel of collagen following incubation with different concentrations of ribose. The dotted lines are for visual reference of mobility changes. All samples had been incubated with ribose for 350 days.

Using SDS-PAGE we observed a monotonic decrease in electrophoretic mobility in both *α* (single *α* chains) and *β* (two cross-linked *α* chains) bands of collagen, indicating increased mass of individual *α* chains due to glycation (Fig. 2B). Further, a marked decrease in *α* band intensity with ribose concentration, with a concomitant intensity increase of higher-weight bands between *β* and *γ* (three cross-linked *α* chains) bands, indicated increased cross-linking with higher ribose concentrations. (The uncropped gel can be seen in Fig. S1.) While these results represent ∼one year of incubation, changes in mobility were detectable at shorter times, as short as two weeks (Fig. S2A). Additionally, mobility changes were observed to increase with incubation time (Fig. S2B).

### Collagen’s susceptibility to cleavage by trypsin is increased by glycation

Susceptibility to trypsin digestion was used to probe the effects of AGEs on the stability of collagen’s triple helix [23, 24]. Figure 3A displays the results of the incubation with trypsin, showing lower band intensity in the trypsinized (+) collagen lanes with respect to the non-trypsinized (-) collagen control lanes. This reduction of intensity increases with ribose concentration (and therefore extent of glycation). (The uncropped gel can be seen in Fig. S3.) A reduction of band intensity (seen in both *α* and *β* bands) indicates digestion of collagen by trypsin. Quantitative analyses of this gel provided the percent reduction of integrated band intensity resulting from trypsin cleavage. Band intensities of both *α* and *β* bands of trypsin (+) and the respective control lanes lacking trypsin (-) were compared and showed very similar patterns, indicating that the destabilization was independent of inter-chain crosslinks induced by AGEs. The results clearly demonstrate an increased susceptibility to digestion with increasing glycation (Fig. 3B). Thus, glycation destabilizes collagen at the molecular level.

**FIG. 3.**
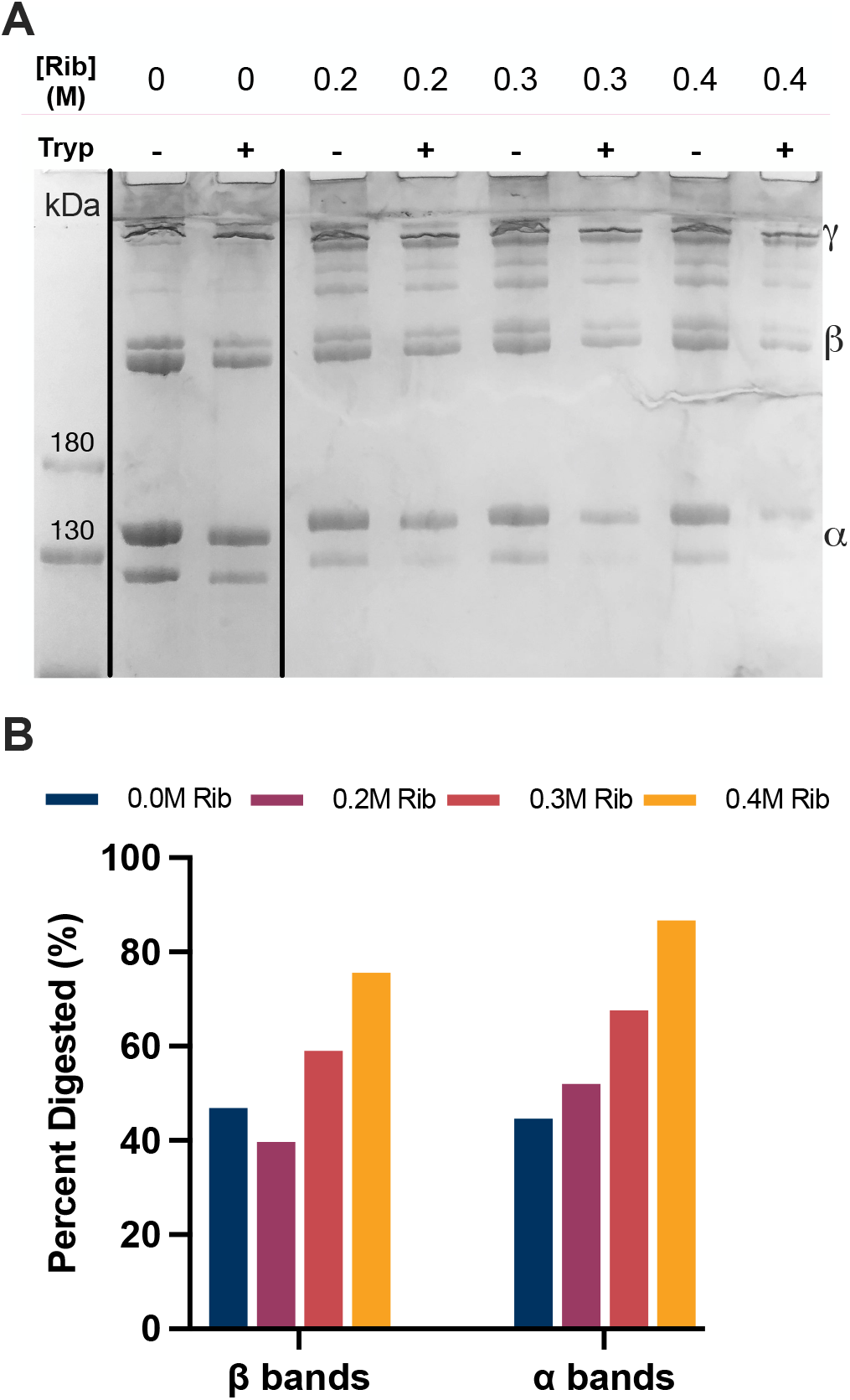
Trypsin digestion of glycated collagen. (A) SDS-PAGE gel of collagen glycated by various concentrations of ribose, followed by trypsin treatment (+) or control (-). Ratios of intensities between adjacent bands (plus and minus trypsin) are depicted in B. (B) Percent digestion of *α* and *β* bands. All samples had been incubated with ribose for 350 days.

### Collagen’s thermal stability is decreased by glycation

To further investigate glycation-induced destabilization, circular dichroism (CD) spectroscopy was used to assess collagen’s structure and thermal stability. Figure 4A shows scaled CD spectra for glycated and nonglycated collagen samples. (Unscaled spectra can be seen in Fig. S4.) Both samples show the characteristic positive peak around 222 nm associated with the right-handed triple helix, and the negative peak around 197 nm corresponding to the left-handed *α* chains. The *R*_pn_ value (ratio of the magnitudes of the positive peak to the negative peak) [25] was essentially constant between the two spectra (0.122 and 0.116 for 0 M and 0.4 M ribose respectively, differing by only ∼ 5%), indicating that collagen’s characteristic structure was preserved. Additionally, the *R*_pn_ of nonglycated collagen was nearly identical (within 0.2%) to that of “fresh” collagen that had not been incubated at room temperature, indicating that room temperature storage did not impact the triple helicity of the collagen.

**FIG. 4.**
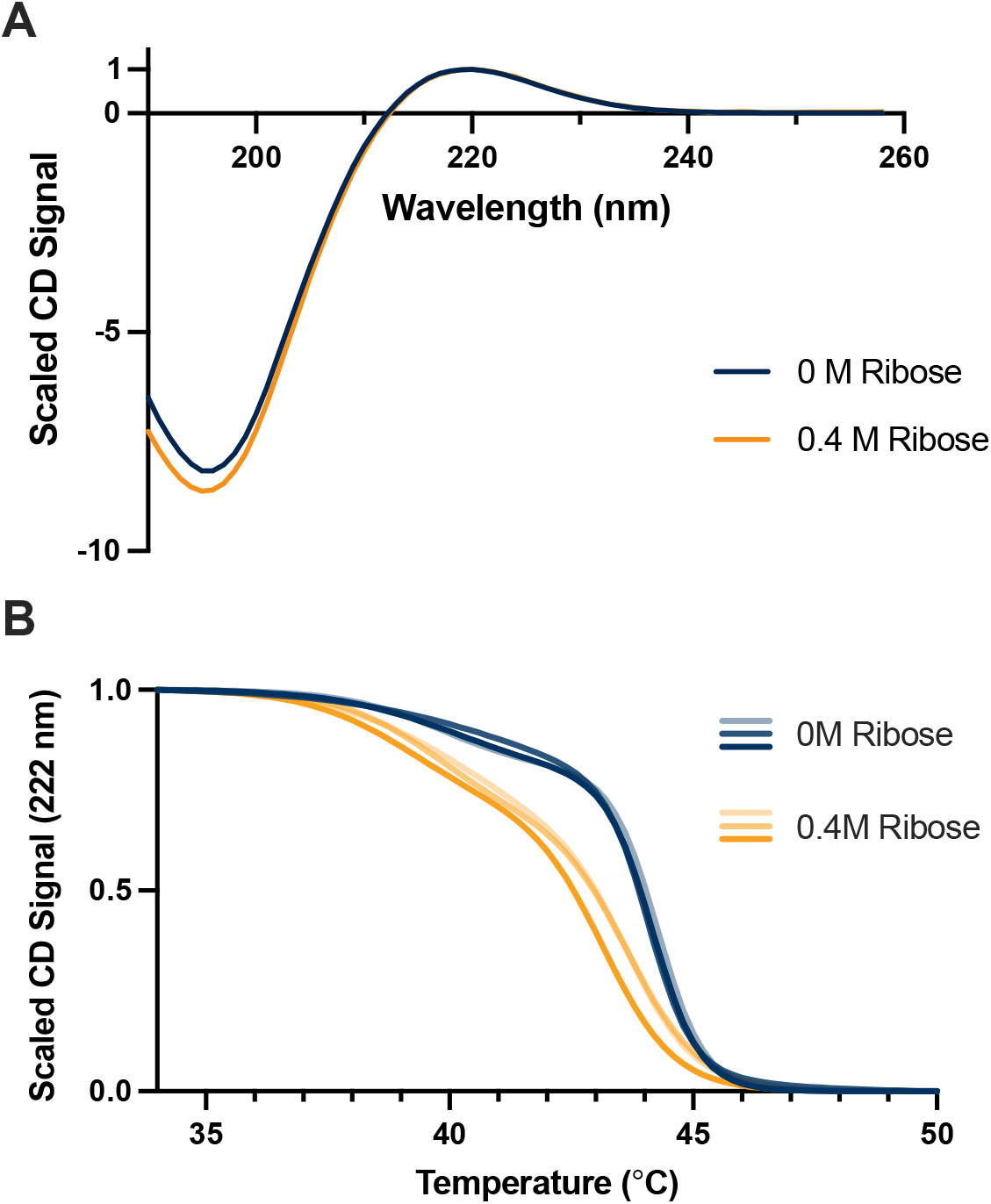
Circular dichroism spectroscopy spectra and melt curves of collagen incubated for 350 days at room temperature with and without ribose. (A) CD spectra of glycated and nonglycated collagen samples, normalized at 222 nm. (B) Thermal denaturation of glycated and nonglycated collagen. A decrease in melting temperature (*T*_m_) due to glycation is apparent. Triplicate measurements are depicted as different shades of yellow and blue for the glycated and nonglycated samples, respectively. Biphasic sigmoid fits to the normalized responses are shown. Full data are shown in Fig. S6.

However, thermal denaturation, monitored via a loss of intensity of the CD signal at 222 nm when heating [26], revealed a significant loss of thermal stability arising from glycation (Fig. 4B). The melting behavior was better described by a two-step, biphasic sigmoid model than a single sigmoid, even for “fresh” collagen that had not been incubated at room temperature (Fig. S5). By determining the mid-point of the transitions – the temperature at which the amplitude of the signal in the biphasic sigmoid fit reduced to half of its initial value – we quantified the apparent melting temperatures: *T*_m_ = (42.83 ± 0.14) °C and (43.94 ± 0.04) °C for glycated and nonglycated samples respectively. These apparent melting temperatures are averaged from three replicate melt curves, with errors representing the standard deviation of *T*_m_ among replicates. Fresh collagen (not incubated at room temperature) had *T*_m_ = (44.5 ± 0.07) °C. It is important to note that these values of T_*m*_ do not represent the actual melting temperature of collagen, which depends on the heating rate [27] (here, 0.5°C/min). Instead, they reveal the differences between samples analyzed under identical conditions, showing that glycated collagen is significantly less thermally stable than its nonglycated control.

### Collagen’s bending stifness is increased by glycation

Building upon these insights that glycated collagen molecules exhibit reduced stability compared to their nonglycated counterparts, we next explored the consequent impact on the mechanical properties of these molecules, focusing on their persistence length, a measure of bending stiffness. Utilizing Atomic Force Microscopy (AFM), we imaged glycated and nonglycated collagen molecules that were deposited then dried on mica substrates [19]. Their contours were traced and parametrized using AutoSmarTrace [20], as shown in Fig. 5A. The persistence length of each sample was obtained from these contours using the Worm-Like-Chain (WLC) model for polymer bending flexibility (Eq. 2, 3). See Fig. S7A for example fits to the WLC using equations 2 and 3. Persistence length quantifies bending stiffness: a longer persistence length means that the chain direction persists for a longer stretch of the molecular backbone, meaning it costs more energy for it to bend and change direction. Conversely, a shorter persistence length means the chain is more flexible.

**FIG. 5.**
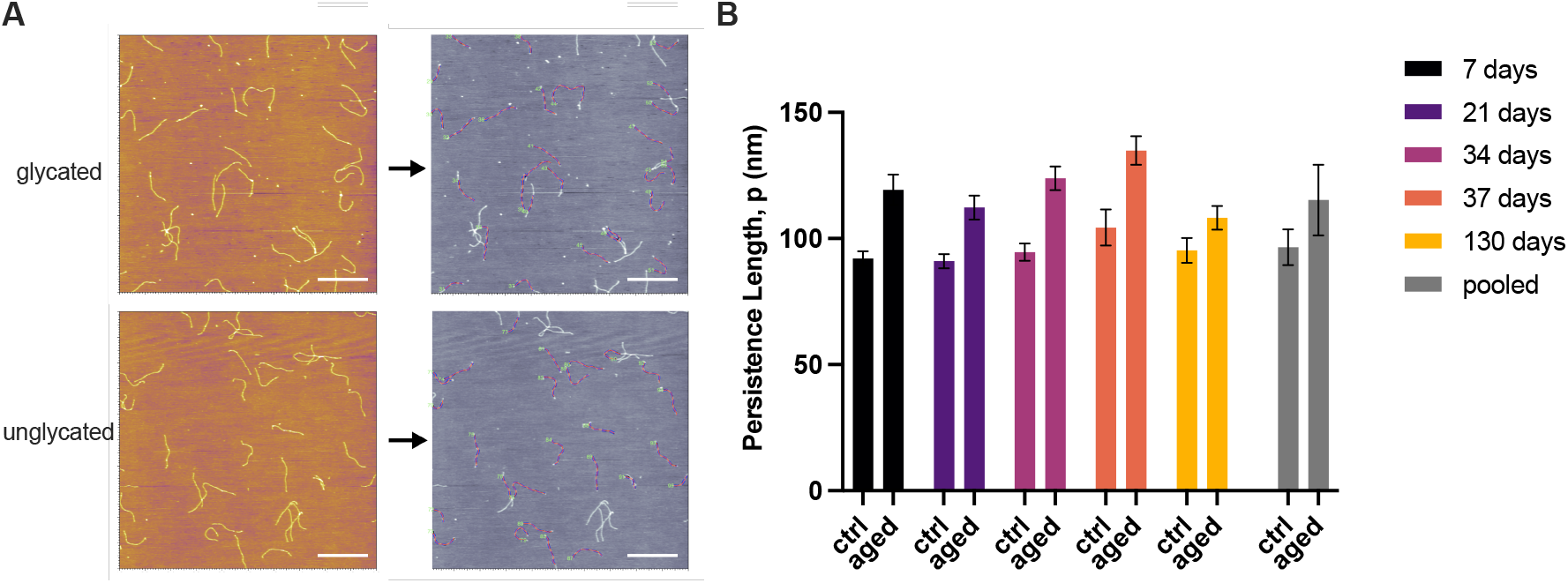
Sample AFM images, their respective contour traces and the determined persistence lengths of various glycated collagen samples and their nonglycated counterparts. (A) Sample AFM images of glycated (0.4 M ribose, 130 days) (top) and control (bottom) (0 M ribose, 130 days) collagen molecules and their resulting traces via AutoSmarTrace. Scale bars are 400 nm. (B) Persistence lengths of collagen samples, encompassing different ribose concentrations and incubation durations. The 7-, 21-, and 34-day datapoints are of the same sample, incubated with 0.4 M ribose, but not dialysed before plating on mica; the 130-day sample was incubated with 0.4 M ribose then dialysed. The 37-day sample was incubated with 1 M ribose, then dialysed. The pooled data results represent the averages and standard deviations of all of these respective control and aged collagen persistence lengths from these different incubation conditions. Errors were generated via bootstrapping. Summary of the data used for this analysis and relevant controls can be seen in Fig. S7B and Table S1.

We found the persistence length of collagen to increase following glycation. This was observed across various changes in glycation protocol (e.g. concentration and duration) (Fig. 5B and Table S1). Surprisingly, we did not observe a trend of increasing persistence length as incubation time increased. The earlier time-point samples (at 7, 21 and 34 days) were not dialysed to remove free ribose (concentration of ∼20 µM upon deposition on mica); however, control experiments show that free ribose does not significantly alter collagen’s persistence length (Fig. S7B). Thus, our results demonstrate that the increase in collagen’s bending stiffness arises from glycation.

### Collagen’s ability to form fibrils is inhibited by glycation

The impact of glycation on collagen’s ability to self-assemble into fibrils was investigated using Differential Interference Contrast (DIC) microscopy. We found DIC microscopy to provide superior contrast to bright-field imaging, which has been used previously to image collagen fibrillar networks [28]. The self-assembly of nonglycated, control collagen into fibrillar matrices is easily discernable with DIC microscopy (Fig. 6A). By contrast, glycated collagen did not undergo self-assembly into fibrils, even after overnight incubation (Fig. 6B).

**FIG. 6.**
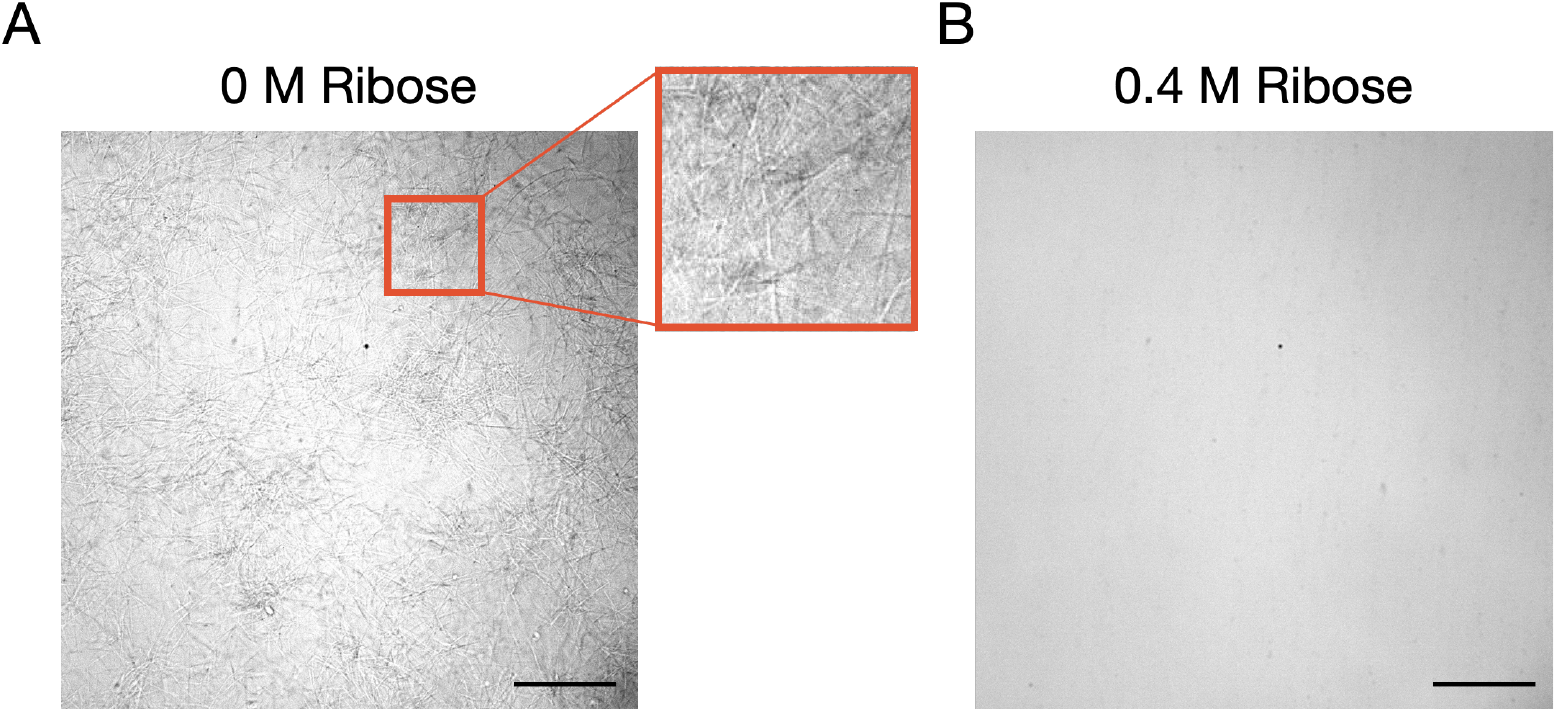
DIC microscopy imaging of collagen fibrillar structures. Collagen was incubated for 42 days with 0 M (A) or 0.4 M (B) ribose, then dialysed to remove the ribose prior to assembly. Collagen was then incubated overnight in PBS, where the control sample formed fibrillar structures, while the glycated collagen did not. Scale bar is 50 µm.

## DISCUSSION

In this study, collagen was incubated with ribose to introduce advanced glycation end-products (AGEs). Although glucose is the more physiologically relevant sugar involved in glycation, we worked with ribose because it has been shown previously to induce glycation effects similar to glucose but at a significantly faster rate [29–31].

Glycation was assessed through two complementary methods: fluorescence measurements at AGE-specific wavelengths, and monitoring changes in electrophoretic mobility with SDS-PAGE. We found that ribose incubation produced a dose-dependent increase in fluorescent AGEs and decrease in mobility of collagen chains in electrophoresis (Fig. 2).

A dose-dependent increase in fluorescence associated with AGEs has been observed in studies of collagen type I (and type IV [32]), incubated with increasing concentrations of reducing sugars such as ribose [32–34], glucose [34, 35], and the sugar derivative and carbohydrate glycolysis byproduct methylglyoxal (MGO) [36]. Similar to prior findings of increasing fluorescence from AGEs as a function of incubation time [37], we also observed a time-dependent increase in AGEs (Fig. S2B). Fluorescence measurements of AGEs have utilized a range of excitation and emission wavelengths (*λ*_ex_/*λ*_em_), including 335/420 [38], 370/425 [37], 360/465 [34] and 320/410 [39], all of which yielded comparable trends to our observations at 328/378 and 370/440 (Fig. 2A). Moreover, our observations of glycation inducing decreased electrophoretic mobility are consistent with prior studies of collagen types I [40] and IV [32]. This validation underscores the reliability and sensitivity of gel mobility analysis in assessing glycation progression. It is worth noting that many AGEs are non-fluorescent [3], which can limit the ability of fluorescence measurements to determine the extent of glycation without the supplemental electrophoretic mobility analysis.

Having established that our collagen samples were chemically modified, we then studied the effect of these AGEs on the structure and stability of collagen’s triple helix. Our circular dichroism (CD) results showed that collagen’s triple-helical structure is maintained at room temperature following glycation, but that its thermal stability decreases (Fig. 4). In our work, we quantify the extent of triple-helical structure by the ratio of the positive peak amplitude at 222 nm to the amplitude of the negative peak at 197 nm, *i*.*e*., by the value of *R*_pn_. *R*_pn_ is independent of concentration, allowing easy comparison between different collagen samples. Our finding of a maintained triple-helical structure aligns with a previous study of glycated type I collagen [41]. In other work, a decrease in signal at 222 nm of type IV collagen following glycation was attributed to a destabilization of the triple helix; this destablization further resulted in a decrease in thermal stability of the AGEd protein [32]. We found a decrease in melting temperature of glycated type I collagen (Fig. 4B), but found that its average structure at room temperature was mostly maintained following glycation (Fig. 4A). Interestingly, previous research found increased thermal stability in tissues extracted from naturally aged rabbit collagen [42], suggesting that protein organization into fibrils may counteract the destabilization of individual collagen molecules. However, varying conditions such as pH, temperature, and physiological context might influence these results.

Somewhat surprisingly, although the triple-helical structure of collagen is little altered by glycation, we did observe a significant enhancement of its susceptibility to digestion by trypsin (Fig. 3). The effects were dose-dependent, with significantly more cleavage observed for samples incubated with higher concentrations of ribose (and posssessing a greater extent of AGEs; Fig. 2). It is possible that glycation does not significantly alter the triple-helical structure overall, but perturbs it enough near trypsin cleavage sites that the individual *α* chains become more susceptible to proteolysis. This picture is supported by the site specificity of glycation and of trypsin proteolysis: trypsin exclusively cleaves C-terminal to lysine and arginine residues [43], which are the primary residues targeted in glycation [3]. Our findings suggest that glycation-related modifications of lysine and arginine residues do not inhibit trypsin access and cleavage; instead, they may promote cleavage. This observation implies that steric effects of glycation modifications may be negligible, or other factors may be influencing proteolytic susceptibility.

While there may be a common destabilizing mechanism for glycation effects on soluble proteins such as collagen and BSA [37], the situation appears to be more complex for collagen fibrils. For instance, glycated collagen I scaffolds have been documented to exhibit increased resistance to degradation by bacterial collagenase [36]. Other investigations have indicated a significant decrease in collagen turnover, specifically degradation by matrix metalloproteinases (MMPs), in glycated tissue, attributed to inhibitory effects of AGEs themselves [44]. These findings demonstrate that AGEs seem to have different effects on collagen’s proteolytic susceptibility depending on its structural level. Intermolecular crosslinks present in glycated fibrils – but not in our molecularly glycated collagen samples – likely play a role in inhibiting proteolytic access to the *α* chains for cleavage.

To further understand the effects of AGEs on collagen at the molecular level, we used AFM to image glycated collagen proteins and quantified their bending stiffness via the persistence length, *p*. We were motivated to study mechanical changes in individual collagen proteins, given previous findings of AGEs stiffening higher-order collagen structures [3, 12, 45, 46]. Our results show that glycation increases the bending stiffness of collagen proteins, increasing the persistence length of collagen by ∼20% (Fig. 5, Table S1). Thus, glycation affects the mechanics of collagen not only by introducing intermolecular crosslinks between neighboring collagen proteins in fibrils and between fibrils [12, 40, 47], increasing their stiffness, but also by stiffening individual collagen proteins.

The effects of glycation on the structure, stability and mechanics of the triple helix of collagen thus are multifaceted: the average triple-helical structure is maintained (Fig. 4A), though it becomes more susceptible to proteolysis (Fig. 3) and less thermally stable (Fig. 4B); and its bending stiffness increases, meaning its flexibility is reduced by glycation.

It is surprising that bending stiffness is not increased by incubation with ribose in a time-nor dose-dependent fashion, in contrast to the extent of glycation and its effects on other aspects of collagen structure and stability. It could be that triple-helix bending is facilitated by an ability of the collagen *α* chains to locally unwind and slide past each other; such motion would be impeded by the crosslinks formed by AGEs between *α* chains of individual proteins. We observe such intramolecular / interchain crosslinks by the decrease in *α*-band intensity of glycated collagens, accompanied by an appearance of higher-weight bands in SDS-PAGE (Fig. 2B). While these increase with ribose concentration, it could be that only a small number of crosslinks is necessary to lock in chain register, such that additional crosslinking does not further increase the bending stiffness. Although the electric charge of collagen may be affected by AGE formation [48], and electrostatics do influence the conformation of collagen proteins on the mica surface used for AFM imaging [19, 49], we previously showed that the dominant effect is on molecular curvature rather than on persistence length [19]. Thus, we do not anticipate a change in collagen’s charge to be responsible for the glycation-dependent increase in measured bending stiffness. We do find some variability in persistence length for different preparations of collagen, even under the same deposition conditions (Table S1). In this study, we have directly compared persistence lengths between collagen incubated with ribose and without, for the same duration of time, so that we control as much as possible for any extrinsic factors influencing our measurements of persistence length (Fig. 5). We have also updated the SmarTrace analysis algorithm to use bootstrapping for estimating uncertainties in values of persistence length. This better captures the variability of conformations among collagen proteins, rather than simply providing an estimated uncertainty in fitting parameters for the entire sampled population. Taken together, we believe these considerations help to provide confidence in the finding that glycation increases the bending stiffness of collagen proteins.

One key attribute of collagen is its ability to self-assemble into higher-order fibrillar structures. We found glycation to quash this ability to assemble. Free ribose in solution was seen to have a strong influence on fibril formation (Fig. S8), slowing, altering and even inhibiting fibril formation with effects seen at ribose concentrations as low as 25 mM. Thus, dialysis was deemed essential to eliminate free ribose before performing any fibril formation experiments on glycated collagen. Even following dialysis, glycated collagen did not assemble into fibrils, while untreated collagen did (Fig. 6). Inhibition of collagen assembly has previously been observed for collagens glycated in molecular form [34], consistent with our observations. We did not test the ability of glycated collagen to undergo self-assembly after very short incubation periods. Previous work found that self-assembly was still possible, though slower, after 24 hours of incubation with a reducing sugar [41]. Clearly, there must be some changeover between no effect (zero time of incubation) to complete inhibition of fibril formation, but elucidating this time dependence was not the focus of our study.

The modification of lysine and arginine side-chains by AGEs will almost certainly alter salt-bridge formation within and between collagen proteins [50–52]. This can disrupt electrostatic interactions important in guiding the register between proteins in fibrils [48]. Indeed, it has previously been reported that even low levels of amine modification in collagen reduce its fibrillogenesis kinetics [53], consistent with our observations. However, it is unclear from this previous study whether the impact was from the chemical modification of amines or from the effects of the tethered cyanine fluorophores. In light of our finding of increased bending rigidity of glycated collagen, it is interesting to speculate that this stiffening may also contribute to inhibiting assembly [54, 55].

While the dominant effects in this study arise from glycation, we have also noted a few examples where extended incubations at room temperature (noticed for samples incubated for 350 days or longer) alter the properties of collagen when compared with “fresh” collagen stored at 4°C. Incubation at room temperature did not significantly affect collagen’s average triple helical structure, though its thermal stability was impacted. Circular dichroism (CD) analysis showed that the non-glycated sample incubated at room temperature had an essentially identical R_*pn*_ value to that of “fresh” collagen. However, the measured *T*_m_ was reduced by ∼0.5°C following extended incubations at room temperature compared with collagen stored at 4°C. This indicates that some thermal destabilization occurred independently of glycation due to room temperature incubation. We also observed a loss of function in nonglycated samples that had sat at room temperature for extended periods of time, namely, these samples were unable to form proper fibrils, suggesting that an intrinsic, time-dependent loss of function unrelated to glycation can also occur.

In this study, glycation was performed at room temperature in an acidic environment (acetic acid, pH ∼3.2) to prevent fibrillogenesis and maintain molecular collagen as described in [33–35]. Although, physiologically, collagens are generally surrounded by neutral pH at ∼37°C, AGE formation has been shown to occur under both acidic and neutral conditions, although quicker at neutral pH [33]. Thus acidic glycation is a suitable method for inducing AGE formation while preserving molecular collagen.

## CONCLUSIONS

In this study we utilized various biochemical techniques combined with single-molecule methods to elucidate the effects of glycation and the accumulation of AGEs on molecular collagen type I. We demonstrated SDS-PAGE and fluorescence measurements to be a reliable methods for tracking glycation, elucidating its time and sugar-concentration dependence. We then found that triple-helical stability is decreased in glycated collagens, which we determined via their increased susceptibility to trypsin proteolysis and decreased melting temperature as seen with CD spectroscopy. Next we used AFM imaging and discovered that this loss of stability was accompanied by a mechanical stiffening or decrease in bending flexibility of individual collagen molecules. Finally we showed evidence of the disruptive impact of glycation on collagen’s self-assembly process, shedding light on the underlying mechanisms driving age-related alterations in collagen structure and function. We believe this is the first demonstration of an effect of AGEs on the mechanics of collagen at the individual protein level.

## ACKNOWLEDGEMENTS

We thank Alaa Al-Shaer for her guidance and assistance with data collection. We are very grateful to Emily Cranston at UBC for providing access to the Asylum Jupiter AFM (funded by CFI). Additionally, we appreciate the advice and useful suggestions from Ryan Bauer, Jesús Serrano-Negrón, Karmen Gill, and Jody Tao in experimental design.

## FUNDING SOURCES

This work was funded by the Natural Sciences and Engineering Research Council of Canada (NSERC) via a Discovery Grant to NRF and an Undergraduate Student Research Award (USRA) to DS. DS was also supported in part by funding from the Canada Summer Jobs program.

## SUPPORTING INFORMATION

Supporting information is available and includes Figures S1-S8 and Table S1.

## Notes

### Competing Interest Statement

The authors have declared no competing interest.

## References

[1] C. Franceschi, M. Capri, D. Monti, S. Giunta, F. Olivieri, F. Sevini, M. P. Panourgia, L. Invidia, L. Celani, M. Scurti, et al., Mechanisms of Ageing and Development 128, 92 (2007).

[2] B. K. Kennedy, S. L. Berger, A. Brunet, J. Campisi, A. M. Cuervo, E. S. Epel, C. Franceschi, G. J. Lithgow, R. I. Morimoto, J. E. Pessin, et al., Cell 159, 709 (2014).

[3] A. J. Bailey, Mechanisms of Ageing and Development 122, 735 (2001).

[4] S. Vasan, X. Zhang, X. Zhang, A. Kapurniotu, J. Bernhagen, S. Teichberg, J. Basgen, D. Wagle, D. Shih, I. Terlecky, et al., Nature 382, 275 (1996).

[5] B. N. Mason, A. Starchenko, R. M. Williams, L. J. Bonassar, and C. A. Reinhart-King, Acta Biomaterialia 9, 4635 (2013).

[6] K. C. Tan, W.-S. Chow, V. H. Ai, C. Metz, R. Bucala, and K. S. Lam, Diabetes Care 25, 1055 (2002).

[7] A. Rojas, C. Añazco, I. González, and P. Araya, Carcinogenesis 39, 515 (2018).

[8] H. Dandia, K. Makkad, and P. Tayalia, Biomaterials Science 7, 3480 (2019).

[9] H. Kuniyasu, N. Oue, A. Wakikawa, H. Shigeishi, N. Matsutani, K. Kuraoka, R. Ito, H. Yokozaki, and W. Yasui, The Journal of Pathology 196, 163 (2002).

[10] R. Paul and A. Bailey, The International Journal of Biochemistry & Cell Biology 28, 1297 (1996).

[11] V. M. Monnier, G. T. Mustata, K. L. Biemel, O. Reihl, M. O. Lederer, D. Zhenyu, and D. R. Sell, Annals of the New York Academy of Sciences 1043, 533 (2005).

[12] A. Gautieri, F. S. Passini, U. Silván, M. Guizar-Sicairos, G. Carimati, P. Volpi, M. Moretti, H. Schoenhuber, A. Redaelli, M. Berli, et al., Matrix Biology 59, 95 (2017).

[13] M. W. Kirkness, K. Lehmann, and N. R. Forde, Current Opinion in Chemical Biology 53, 98 (2019).

[14] C. Yang, S. Mosler, H. Rui, B. Baetge, H. Notbohm, and P. K. Müller, Matrix Biology 14, 643 (1995).

[15] S. E. Hormel and D. R. Eyre, Biochimica et Biophysica Acta (BBA)-Protein Structure and Molecular Enzymology 1078, 243 (1991).

[16] R. R. Kohn and E. Rollerson, Journal of Gerontology 15, 10 (1960).

[17] M. Bellmunt, M. Portero, R. Pamplona, M. Muntaner, and J. Prat, Lung 173, 177 (1995).

[18] D. R. Sell and V. M. Monnier, Journal of Biological Chemistry 264, 21597 (1989).

[19] N. Rezaei, A. Lyons, and N. R. Forde, Biophysical Journal 115, 1457 (2018).

[20] M. Schneider, A. Al-Shaer, and N. R. Forde, Biophysical Journal 120, 2599 (2021).

[21] M. Shayegan and N. R. Forde, PLOS ONE 8, e70590 (2013).

[22] K. E. Kadler, D. F. Holmes, J. A. Trotter, and J. A. Chapman, Biochemical Journal 316, 1 (1996).

[23] P. Bruckner and D. J. Prockop, Analytical Biochemistry 110, 360 (1981).

[24] M. W. Kirkness and N. Forde, Biophysical Journal 114, 570 (2018).

[25] Y. Feng, G. Melacini, J. P. Taulane, and M. Goodman, Journal of the American Chemical Society 118, 10351 (1996).

[26] N. J. Greenfield, Nature Protocols 1, 2876 (2006).

[27] E. Leikina, M. Mertts, N. Kuznetsova, and S. Leikin, Proceedings of the National Academy of Sciences 99, 1314 (2002).

[28] A. Kowalewski and N. R. Forde, PLOS ONE 19, e0292298 (2024).

[29] R. R. Kohn, A. Cerami, and V. M. Monnier, Diabetes 33, 57 (1984).

[30] T. Girton, T. Oegema, and R. T. Tranquillo, Journal of Biomedical Materials Research 46, 87 (1999).

[31] M. Brownlee, Diabetes care 15, 1835 (1992).

[32] H.-M. Raabe, H. Molsen, S.-M. Mlinaric, Y. Açil, G. H. Sinnecker, H. Notbohm, K. Kruse, and P. K. Müller, Biochemical Journal 319, 699 (1996).

[33] R. Roy, A. Boskey, and L. J. Bonassar, Journal of Biomedical Materials Research Part A 93, 843 (2010).

[34] N. Diamantides, L. Slyker, S. Martin, M. R. Rodriguez, and L. J. Bonassar, Journal of Biomedical Materials Research Part A 110, 1953 (2022).

[35] M. M. Rowe, W. Wang, P. V. Taufalele, and C. A. Reinhart-King, Soft Matter 18, 8504 (2022).

[36] M. Vaez, M. Asgari, L. Hirvonen, G. Bakir, E. Khattignavong, M. Ezzo, S. Aguayo, C. M. Schuh, K. Gough, and L. Bozec, Acta Biomaterialia 155, 182 (2023).

[37] Y. Wei, L. Chen, J. Chen, L. Ge, and R. Q. He, BMC Cell Biology 10, 1 (2009).

[38] S. Patel, F. Yue, S. Saw, R. Foguth, J. Cannon, J. Shannahan, S. Kuang, A. Sabbaghi, and C. Carroll, Scientific Reports 9, 12614 (2019).

[39] L. Chen, Y. Wei, X. Wang, and R. He, PLOS ONE 5, e9052 (2010).

[40] S.-F. Tian, S. Toda, H. Higashino, and S. Matsumura, The Journal of Biochemistry 120, 1153 (1996).

[41] K. L. Reigle, G. Di Lullo, K. R. Turner, J. A. Last, I. Chervoneva, D. E. Birk, J. L. Funderburgh, E. Elrod, M. W. Germann, C. Surber, et al., Journal of Cellular Biochemistry 104, 1684 (2008).

[42] H. Trebacz, A. Szczesna, and M. Arczewska, Journal of Thermal Analysis and Calorimetry 134, 1903 (2018).

[43] J. V. Olsen, S.-E. Ong, and M. Mann, Molecular & Cellular Proteomics 3, 608 (2004).

[44] J. DeGroot, N. Verzijl, M. Budde, J. W. Bijlsma, F. P. Lafeber, and J. M. TeKoppele, Experimental Cell Research 266, 303 (2001).

[45] O. G. Andriotis, K. Elsayad, D. E. Smart, M. Nalbach, D. E. Davies, and P. J. Thurner, Biomedical optics express 10, 1841 (2019).

[46] R.-Z. Tang and X.-Q. Liu, Materials Today Bio 19, 100607 (2023).

[47] A. Gautieri, A. Redaelli, M. J. Buehler, and S. Vesentini, Matrix Biology 34, 89 (2014).

[48] S. Bansode, U. Bashtanova, R. Li, J. Clark, K. H. Müller, A. Puszkarska, I. Goldberga, H. H. Chetwood, D. G. Reid, L. J. Colwell, et al., Scientific Reports 10, 3397 (2020).

[49] H. H. Lovelady, S. Shashidhara, and W. G. Matthews, Biopolymers 101, 329 (2014).

[50] J.-D. Malcor, N. Ferruz, S. Romero-Romero, S. Dhingra, V. Sagar, and A. A. Jalan, bioRxiv 10.1101/2024.02.24.581883 (2024).

[51] T. Gurry, P. S. Nerenberg, and C. M. Stultz, Biophysical Journal 98, 2634 (2010).

[52] Y. Li and E. P. Douglas, Colloids and Surfaces B: Biointerfaces 112, 42 (2013).

[53] S. Han, D. J. McBride, W. Losert, and S. Leikin, Journal of Molecular Biology 383, 122 (2008).

[54] S. Perret, C. Merle, S. Bernocco, P. Berland, R. Garrone, D. Hulmes, M. Theisen, and F. Ruggiero, Journal of Biological Chemistry 276, 43693 (2001).

[55] F. H. Silver, J. W. Freeman, and G. P. Seehra, Journal of Biomechanics 36, 1529 (2003).

