## Supporting information for "AGEing of Collagen: The Effects of Glycation on Collagen’s Stability, Mechanics and Assembly"

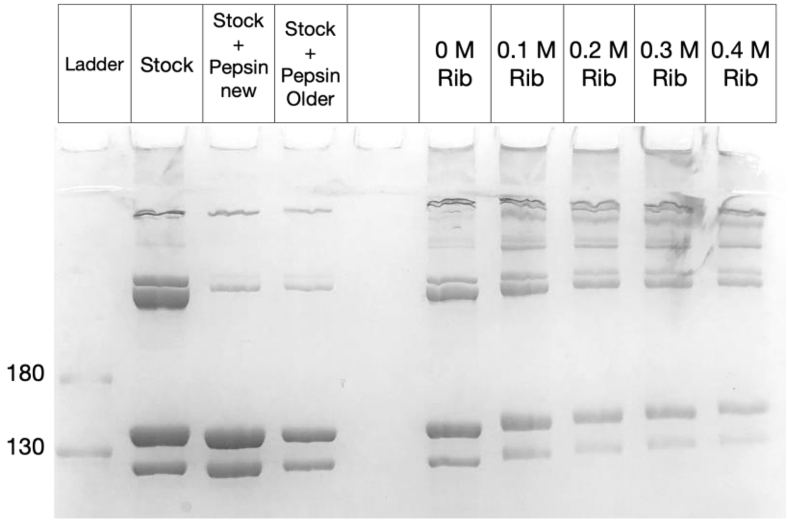


Figure S1. Full gel of glycated collagen samples (Fig. 2 B), cropped in main text to eliminate the pepsin-treated lanes, which are not relevant to the current work. Ladder molecular weights are in units of kDa.

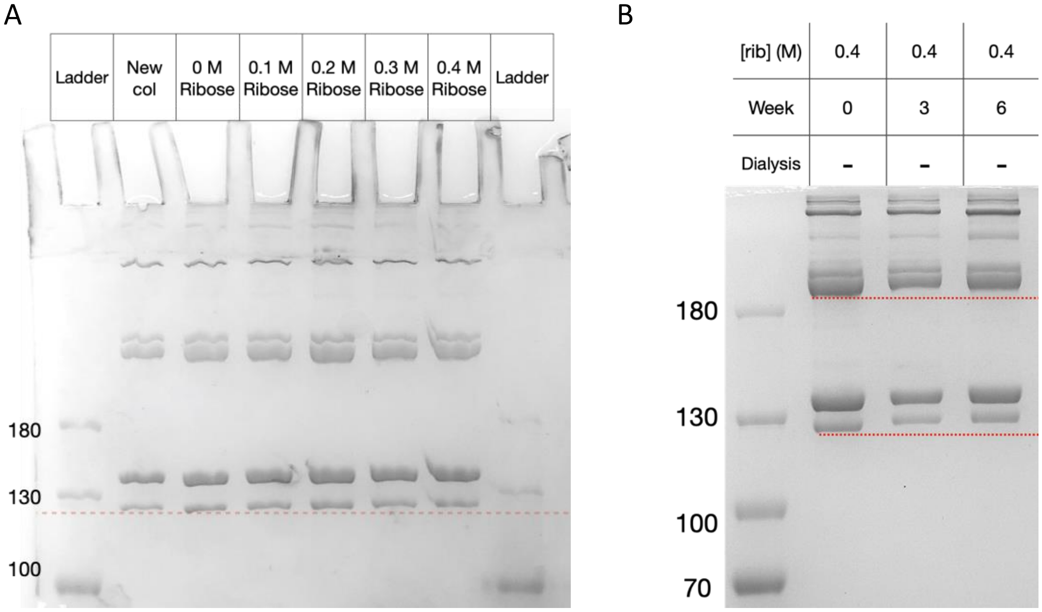


Figure S2. (A) SDS-PAGE gel of collagen incubated with ribose for 14 days exhibits moderate changes in mobility. “New col” is collagen that had been stored at 4°C rather than incubated at room temperature. Comparison with bands in the 0 M ribose lane shows that collagen is not degraded by incubation at room temperature during the glycation protocol. (B) Collagen incubated with a constant concentration of ribose for varying durations shows changes in mobility. The dotted lines are for visual reference of mobility changes. Ladder molecular weights are in units of kDa.

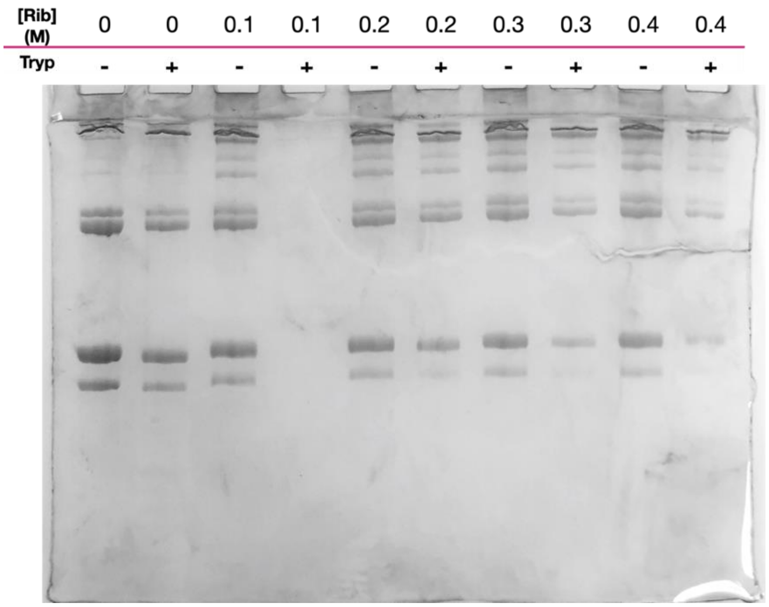


Figure S3. Full gel of the trypsin digest experiments (Fig. 3A), cropped in the manuscript to eliminate lanes 3 and 4. (Lane 4 had a loading error.)

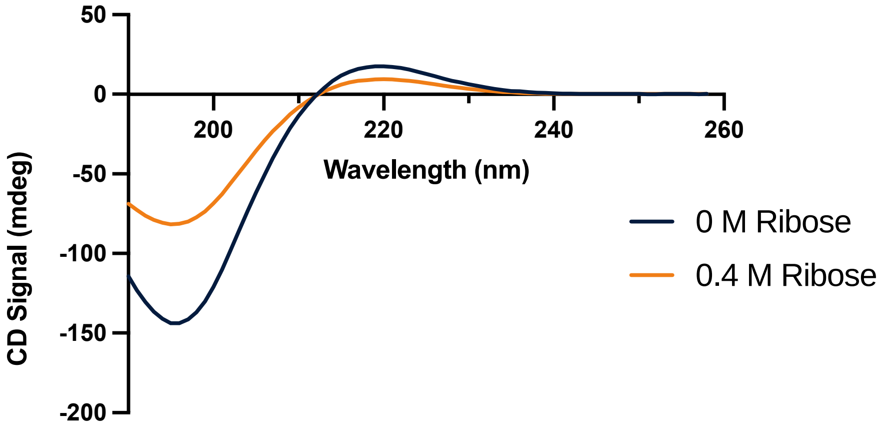


Figure S4. CD spectra of glycated and nonglycated collagen, measured at room temperature. These measurements were rescaled to account for concentration differences, and are replotted in Fig. 4 of the manuscript.

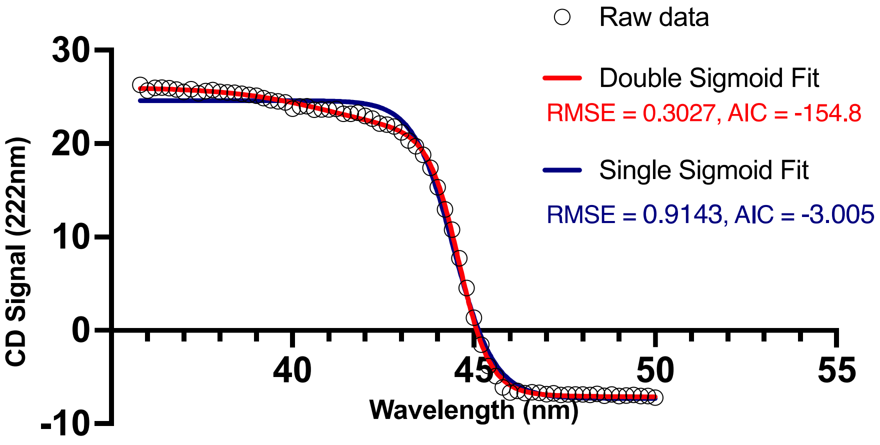


Figure S5. Comparison of single sigmoid and biphasic sigmoid fits to the melt curve of "fresh" collagen (from 4°C storage). While both models can describe the data, statistical parameters (RMSE, AIC) indicate that the biphasic sigmoid is a better fit. Melting temperatures were found to be $T_m$ = 44.5°C and 44.6°C using the single and biphasic sigmoid fits, respectively.

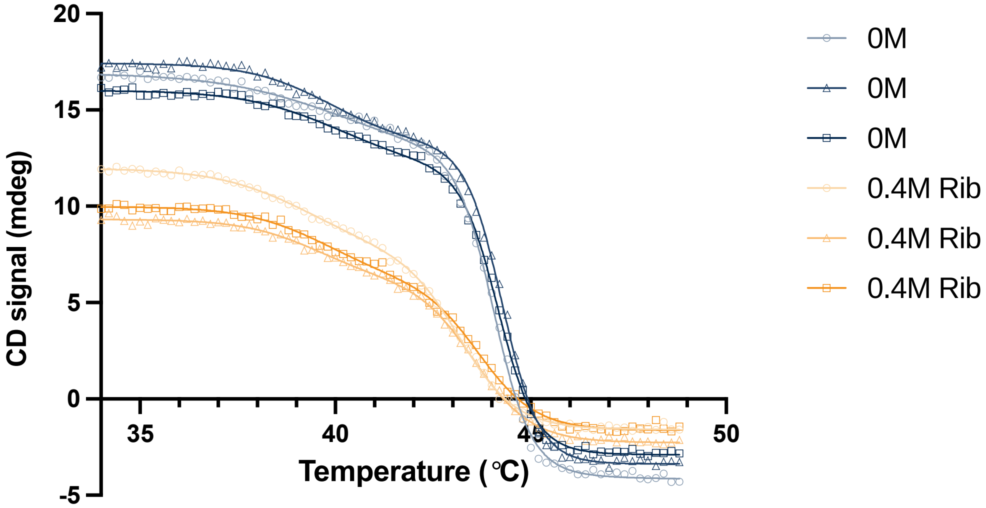


Figure S6. CD melt curves of glycated and nonglycated collagen, with their biphasic sigmoid fits. Data were recorded at 222 nm, with a scanning rate of 0.5°C/min. The biphasic-sigmoid model fits the data very well, with reduced $\chi^2$ values between 0.99 and 1.00 for each fit. Melting temperatures were found to be (42.83 ± 0.14)°C and (43.94 ± 0.04)°C for glycated and nonglycated samples respectively.

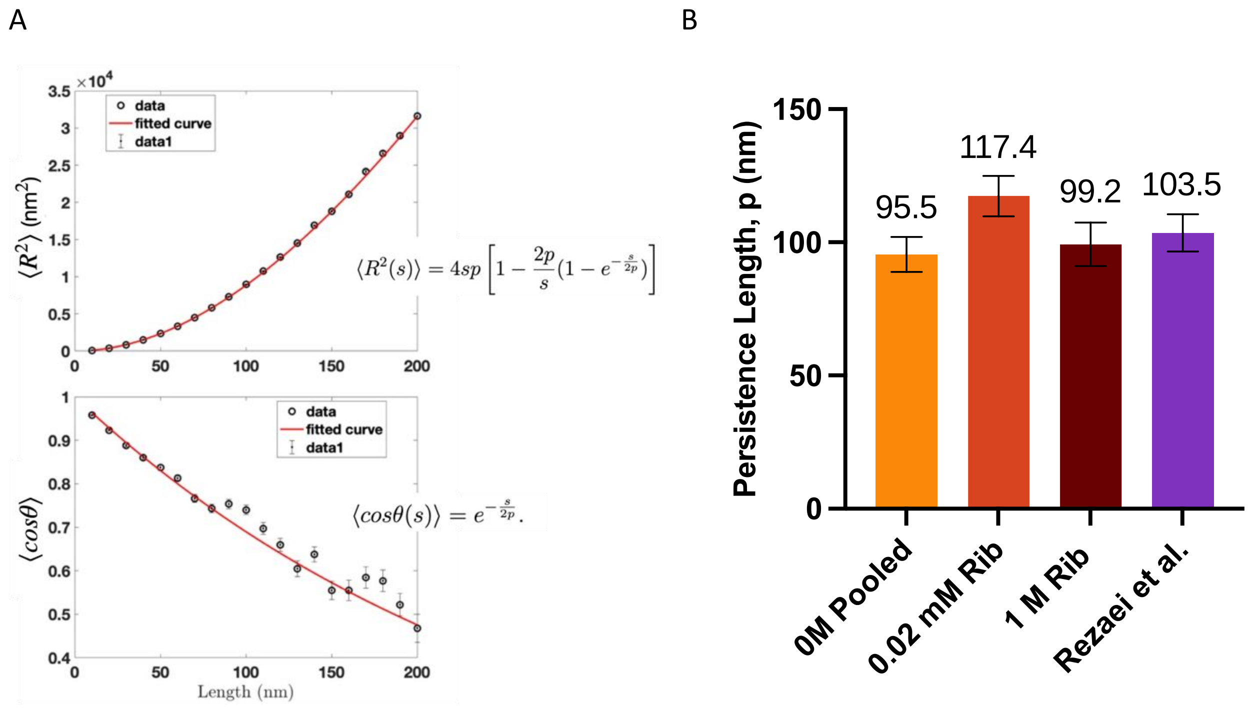


Figure S7. (A) Example fits to the respective WLC equations, for collagen incubated with 1 M ribose for 37 days. (B) The persistence length of non-glycated collagen in the presence of different concentrations of ribose was determined to evaluate the impact of ribose in solution during AFM imaging and the necessity of dialysis. The concentration of 0.02 mM ribose corresponds to the equivalent concentration present during plating for the 7, 21, and 34-day datapoints. Although the persistence length of collagen in the presence of 0.02 mM ribose is higher than the pooled result in the absence of ribose, there is no significant change of persistence length between 0 M and the much higher concentration of 1 M ribose. This indicates that ribose on its own does not strongly impact collagen flexibility. Errors represent standard deviations obtained from bootstrapping iterations, except for the previous results from Rezaei *et al.*, *Biophysical Journal* 2018, for which the error represents the average 95% confidence interval for fits to both mean-squared end-to-end distance and the tangent vector correlation equations.

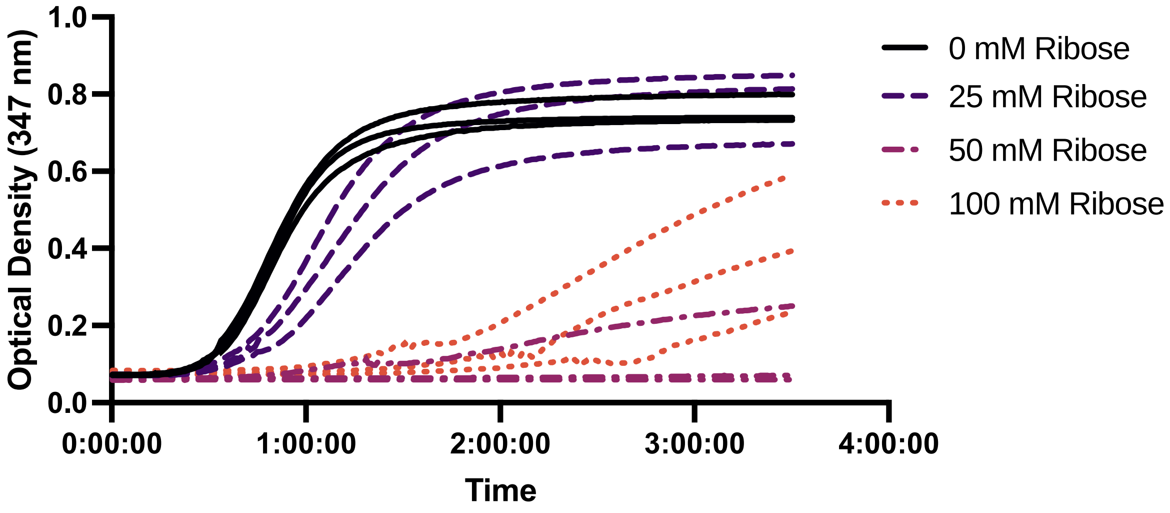


Figure S8. Free ribose interferes with fibril formation in a concentration-dependent manner. Even 25 mM free ribose increases the nucleation phase and decreases the growth rate of fibrils. These experiments utilized fresh collagen, so as to test the effect of different concentrations of free ribose (legend at right) on fibril formation. Turbidity measurements were made in a microplate reader.

| **Incubation time (days)** | **Ribose (M)** | **Dialyzed** | **N chains** | **Average traced chain length (nm)** | **Persistence Length, p (nm)** | **Std. Dev. p (nm)** |
|---|---|---|---|---|---|---|
| 7 | 0.4 | no | 182 | 238.3 | 119.3 | 6.0 |
| 7 | 0 | no | 368 | 235.7 | 92.1 | 2.9 |
| 21 | 0.4 | no | 220 | 236.5 | 112.3 | 4.7 |
| 21 | 0 | no | 299 | 236.2 | 91.0 | 2.8 |
| 34 | 0.4 | no | 166 | 232.6 | 123.9 | 4.7 |
| 34 | 0 | no | 84 | 201.9 | 94.6 | 3.4 |
| 37 | 1 | yes | 138 | 243.4 | 134.9 | 5.7 |
| 37 | 0 | yes | 47 | 206.4 | 104.4 | 7.1 |
| 130 | 0.4 | yes | 277 | 241.9 | 108.2 | 4.7 |
| 130 | 0 | yes | 159 | 235.9 | 95.3 | 4.9 |

Table S1. Details of samples used for AFM analysis. The shorter incubation results were sampled from one preparation following 7, 21 and 34 days of incubation. The 37- and 130-day samples were dialyzed from different preparations at those timepoints, prior to dilution, plating and imaging. Reported persistence length averages and standard deviations are determined as described in the Methods.
